# Zinc Differentially Modulates Tau Aggregation, Fibril Morphology, and Prion-like Seeding in a Construct-Dependent Manner

**DOI:** 10.64898/2026.07.01.735859

**Authors:** Emma L. Poirier, Alyssa Stainton, Oskar Simon, Suhana S. Mittal, Ana B. Varona Ortiz, Sally A. Kim, Jennifer N. Rauch

## Abstract

The role of tau fibril structure in seeding and propagation of aggregation remains a central unresolved question in tauopathy biology. While non-proteinaceous cofactors are increasingly observed in patient-derived tau filaments, whether they actively determine fibril structure and function is not well understood. Here, we show that zinc, a divalent cation dysregulated in Alzheimer’s disease (AD), can drive fundamentally different aggregation and seeding outcomes depending on tau sequence context. Using heparin-free conditions, we compared full-length 2N4R tau (residues 1-441) with an AD-tau fragment (residues 304-380) corresponding to the ordered fibril core. Strikingly, Zn^2+^ exerted opposite effects on these constructs: it accelerated aggregation, increased fibril length, and enhanced cellular seeding for AD-tau, while slowing aggregation, shortening fibrils, and suppressing seeding for full-length tau. These findings demonstrate that cofactor effects are not intrinsic properties of the cofactor itself, but emerge from its interplay with tau sequence and conformational constraints. More broadly, our results support a model in which small-molecule cofactors act as active architects of fibril structure and function, suggesting that chemically distinct environments could generate structurally and biologically distinct tau strains in disease.

## INTRODUCTION

Microtubule-associated protein tau (tau) is an intrinsically disordered protein that is highly enriched in neuronal axons^1^. In diseases such as Alzheimer’s Disease (AD), tau aggregates into neurofibrillary tangles (NFTs), a pathological hallmark of the disease^2^. Tau pathology spreads throughout the brain in a stereotypical pattern^3^, consistent with the leading model that tau pathology spread occurs via a self-propagating mechanism. In this proposed mechanism, aggregated tau templates monomeric tau to adopt a similar aggregation state, a phenomenon known as “seeding”^4,5^. This templated propagation underlies tau’s classification as a “prion-like” protein^6,7^. Understanding the principles governing tau aggregation and seeding is essential for understanding AD progression and developing therapeutic interventions.

Recent cryo-electron microscopy (cryo-EM) studies of AD and other tauopathies have revealed that patients with different tauopathies have distinct tau fibril core structures, while patients with the same disease share a common fibril structure^8,9^. This raises the possibility that unique fibril structures may underlie distinct patterns of disease progression, though the mechanisms linking the two remain unknown. Notably, the cryo-EM studies have also revealed the presence of different unknown densities, varying in size and location across the different folds^8,9^, which may hold a clue to this relationship. Potential candidates for these densities include post-translational modifications, lipids, hydrophobic moieties, or ions, depending on the characteristics of the neighboring residues^10^.

Ions, such as the trace metal zinc (Zn^2+^), have been shown to be dysregulated in the brains of AD patients^11–14^. Zinc concentrations are elevated in AD patient brains compared to control brains, and higher zinc levels correlate with more severe dementia symptoms^11,12^. The dysregulation of zinc is notable, as zinc binds readily to proteins with roughly 10% of the human proteome capable of zinc binding^15^, including tau^16–22^. Zinc is highly abundant in the brain, with average concentrations around 150μM, and at the synapse concentrations can reach up to 1mM^23,24^. Zinc binds directly to tau within its microtubule binding region^16,17^. Functionally, zinc accelerates the *in vitro* aggregation kinetics of both tau-fragments and full-length tau in the presence of heparin, a common inducer of tau aggregation^17–22^. However, comparing aggregation studies across research groups is difficult, as conditions such as shaking, buffer composition, and the tau construct used vary considerably between studies. Moreover, it remains unknown how zinc impacts the seeding potential of tau fibrils. Understanding how zinc shapes tau aggregation, fibril structure, and seeding behavior may therefore clarify not only zinc’s specific role in tauopathy, but may also elucidate broader principles by which co-factors influence fibril structure and how fibril structure, in turn, governs seeding potential.

This study investigates the role of the co-factor zinc (Zn^2+^) in the formation of *in vitro* tau fibrils, the morphology of the resulting fibrils, and their seeding potential in cells. We conducted experiments using two different tau constructs, 2N4R-tau (residues 1-441) the longest brain isoform of tau, and an AD-tau construct (residues 304-380) consisting of the minimal core of the AD tau fibril fold^8^. Strikingly, zinc had opposing effects depending on the construct used. With the shorter AD-tau construct, zinc increased aggregation kinetics, produced slightly longer, narrower fibrils, and increased cellular seeding potential. With the longer 2N4R-tau construct, zinc instead decreased aggregation kinetics, produced shorter, narrower fibrils, and decreased cellular seeding potential. Together, these findings link fibril aggregation, structure, and seeding potential as interdependent properties rather than separate outcomes and highlight that the choice of tau construct used for *in vitro* studies can fundamentally shape the resulting biology.

## RESULTS

### The addition of Zn^2+^ during aggregation alters tau aggregation kinetics but does not impact the amount of fibril formed

To investigate the impact of zinc on tau fibril morphology and seeding potential, we began by generating tau fibrils in the presence and absence of added zinc. Fragments of tau surrounding the microtubule binding region are known to have enhanced aggregation kinetics, thus for our studies we chose to compare two constructs of tau; full-length (“2N4R-tau”) and a shortened fragment comprising just the residues in the ordered AD fibril core (“AD-tau”; residues 304-380) (**Figure 1A**). Both constructs were purified from *E. coli* and aggregation was initiated by incubating tau protein in a shaking plate-reader with or without added ZnSO_4_ using the fluorescent dye thioflavin T (ThT) to track aggregation^25^. ThT aggregation assays with AD-tau demonstrated that added ZnSO_4_ (“(+) Zn^2+^", green) speeds up aggregation kinetics compared to the apo condition without added ZnSO_4_ (“Apo”, grey; **Figure 1B**). Consistent with prior literature^26^, day-to-day variability of tau aggregation kinetics were high (**Supplementary Figure 1A**). Despite this variability, ZnSO_4_ consistently shortened aggregation time by an average of 4h across six independent trials (**Figure 1C**).

**Figure 1.**
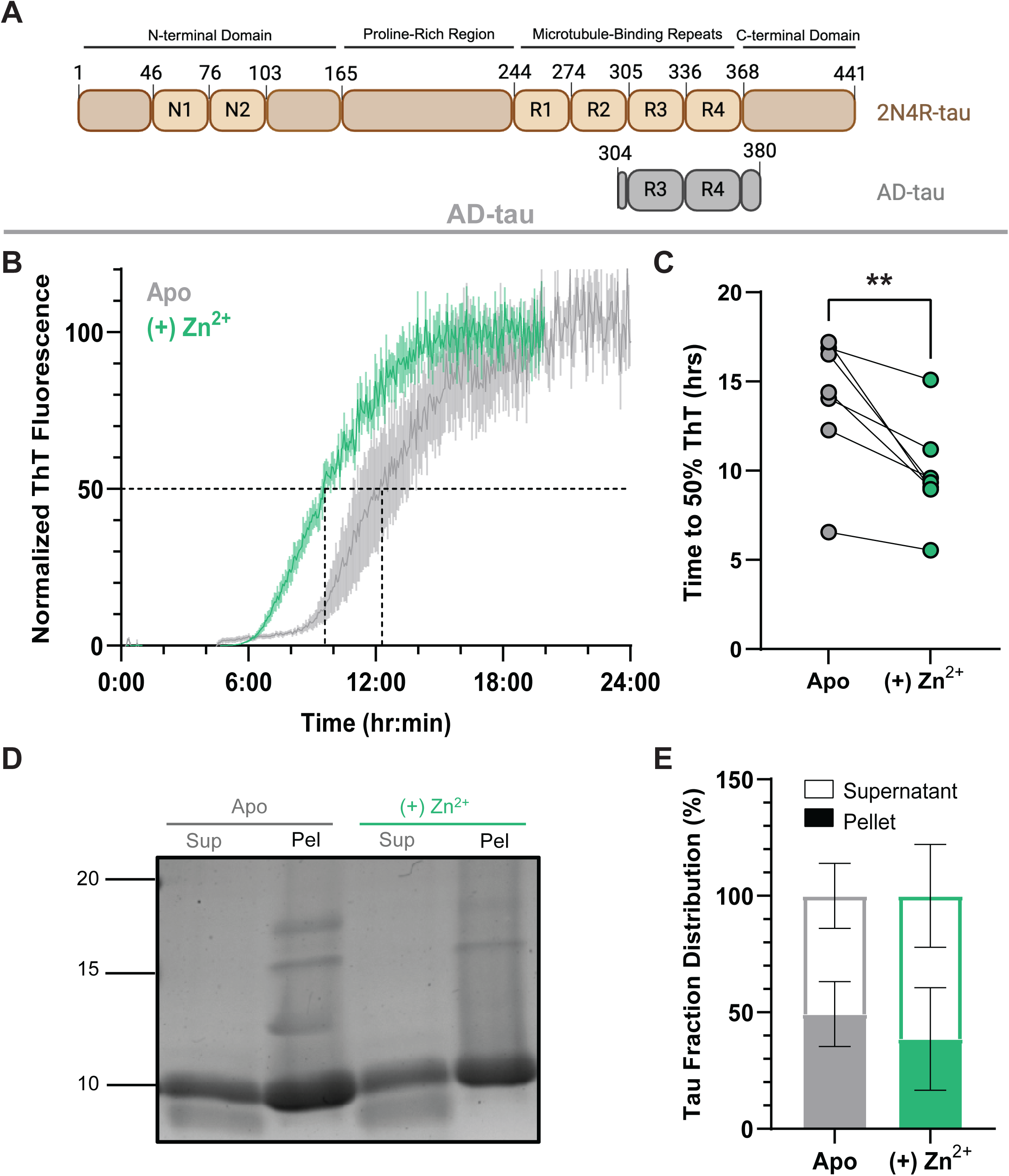
Added Zn^2+^ during AD-tau aggregation increases kinetics but does not impact the amount of fibril formed. A) Schematic representation of the 2N4R-tau and AD-tau constructs used throughout this study. B) Representative AD-tau ThT aggregation assay under single orbital shaking with 50µM tau alone (“Apo,” grey) or with 1.3mM ZnSO_4_ (“(+) Zn^2+^,” green). n=3 technical replicates, each normalized individually. Mean ± SEM plotted. Additional biological replicates (for total N=7) are shown in Supplemental Figure 1A. C) Y_50_ (time in hours to reach 50% of maximum ThT fluorescence, calculated from Gompertz curve fitting) plotted for 7 biological replicates, each with at least 3 technical replicates. Each point represents the mean of one biological replicate. **p = 0.0076, paired t-test. D) Representative SDS-PAGE of supernatant (Sup) and pellet (Pel) fractions after ultracentrifugation of AD-tau aggregation. E) Pellet fraction as a percentage of total tau (Sup + Pel), quantified by densitometry, from 5 biological replicates. Mean ± SEM plotted. No significant difference using a paired t-test.

Following aggregation, fibrils were isolated by ultracentrifugation to separate the supernatant fraction (“Sup”) containing non-aggregated tau from the pellet fraction (“Pel”) containing fibrillized tau. Samples for each were run on an SDS-PAGE gel and quantified with ImageJ to compare total fibrillization efficiency (**Figure 1D**). We found no significant difference in the amount of tau in the pellet fraction generated by the apo and (+) Zn^2+^ conditions (**Figure 1E**) implying that while aggregation kinetics are accelerated the (+) Zn^2+^ condition does not lead to more fibril being made.

Unexpectedly, ThT aggregation assays with the 2N4R-tau construct revealed an opposite trend. With 2N4R-tau, added ZnSO_4_ (“(+) Zn^2+^", blue) slowed aggregation kinetics compared to the apo condition without added ZnSO_4_ (“Apo”, brown; **Figure 2A**). Similar to AD-tau, individual days showed high variability (**Supplementary Figure 1B**). However, we still observed a consistent lengthening of the time to reach 50% aggregation by an average of 23 h across six independent trials (**Figure 2B**). Reports have suggested that shaking parameters can greatly impact tau aggregation efficiency^27,28^, thus we also tested whether performing our assay using single or double-orbital shaking could impact our results. Regardless of shaking condition, we saw the same trend (slowing of aggregation in the presence of ZnSO_4_). Additionally, we observed no significant difference in total aggregation times between single and double orbital shaking for either condition (**Supplemental Figure 1C,D**).

**Figure 2.**
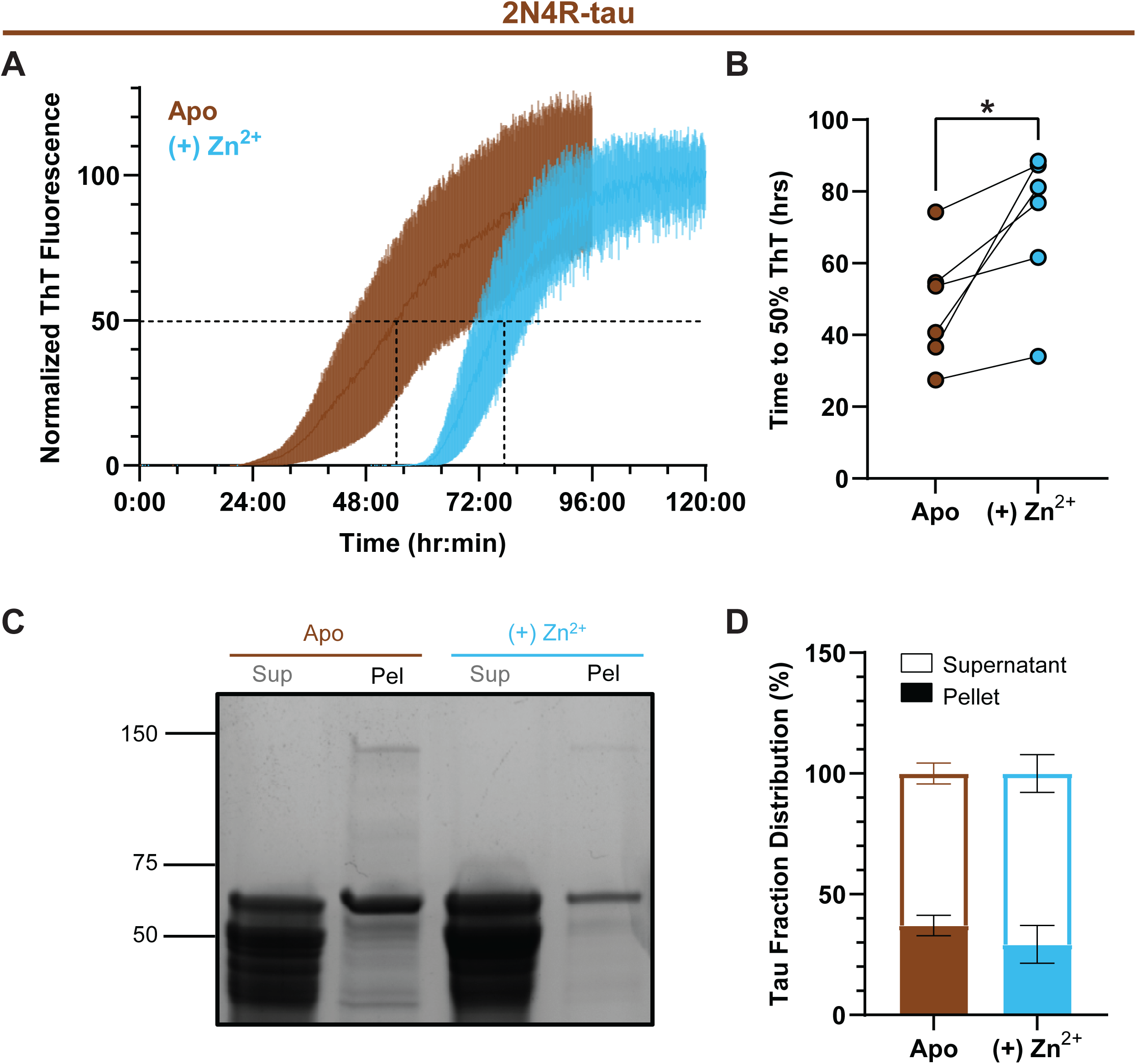
Added Zn^2+^ during 2N4R-tau aggregation decreases kinetics but does not impact the amount of fibril formed. A) Representative 2N4R-tau ThT aggregation assay under double orbital shaking with 50µM tau alone (“Apo,” brown) or with 1.3mM ZnSO_4_ (“(+) Zn^2+^,” blue). n=3 technical replicates, each normalized individually. Mean ± SEM plotted. Additional biological replicates (for total N=6) are shown in Supplemental Figure 1B. B) Y_50_ (time in hours to reach 50% of maximum ThT fluorescence, calculated from Gompertz curve fitting) plotted for 6 biological replicates, each with at least 3 technical replicates. Each point represents the mean of one biological replicate. *p = 0.0260, paired t-test. C) Representative SDS-PAGE of supernatant (Sup) and pellet (Pel) fractions after ultracentrifugation of 2N4R-tau aggregation. D) Pellet fraction as a percentage of total tau (Sup + Pel), quantified by densitometry, from 4 biological replicates. Mean ± SEM plotted. No significant difference using a paired t-test.

To assess total fibril amount, fibrils were isolated by ultracentrifugation and run on an SDS-PAGE gel (**Figure 2C**). Similar to our results with AD-tau, there was no significant difference between apo and (+) Zn^2+^ conditions in the amount of tau in the pellet fraction (**Figure 2D**).

Our aggregation assays were performed in the absence of external inducers, such as heparin, which are known to accelerate aggregation but produce heterogenous filaments that do not resemble brain-derived structures^29^. As prior reports, assessing the impact of zinc on tau fibrillization have all been reported with the use of heparin^17–19,22^, we also examined the effect of zinc on heparin-induced aggregation under our assay conditions. Inclusion of heparin in aggregation assays enhanced aggregation kinetics for both constructs, as expected, but also masked the effect of add Zn^2+^ (**Supplementary Figure 2A-D**). Heparin inclusion also increased the total amount of aggregated tau recovered for both constructs as assessed by SDS-PAGE gels, but similar to our non-heparin assays, total amount of pelleted tau was not influenced by (+) Zn^2+^ conditions (**Supplementary Figure 2E-H**).

### Aggregation with Zn^2+^ changes morphological features of tau fibrils

To assess whether Zn^2+^ impacts fibril morphology, we used negative stain transmission electron microscopy (TEM) to assess fibril length and width. For AD-tau fibrils, the length of fibrils aggregated with zinc were slightly longer than the apo fibrils, with a median length of 555nm and 437nm, respectively (**Figure 3A,B**). The width of the AD-tau fibrils made in the presence of zinc were significantly narrower with a median width of 14.5nm for the (+) Zn^2+^ fibrils vs. 18nm for apo (**Figure 3C**). Contrastingly, for 2N4R-tau fibrils, the (+) Zn^2+^ fibrils were significantly shorter, with a median length of 180nm, compared to the apo fibrils that had a median length of 214nm (**Figure 3D,E**). Similar to the trend observed with AD-tau fibrils, 2N4R-tau (+) Zn^2+^ fibrils were narrower, with a median width of 13nm compared to the apo fibril’s median width of 17nm (**Figure 3F**).

**Figure 3.**
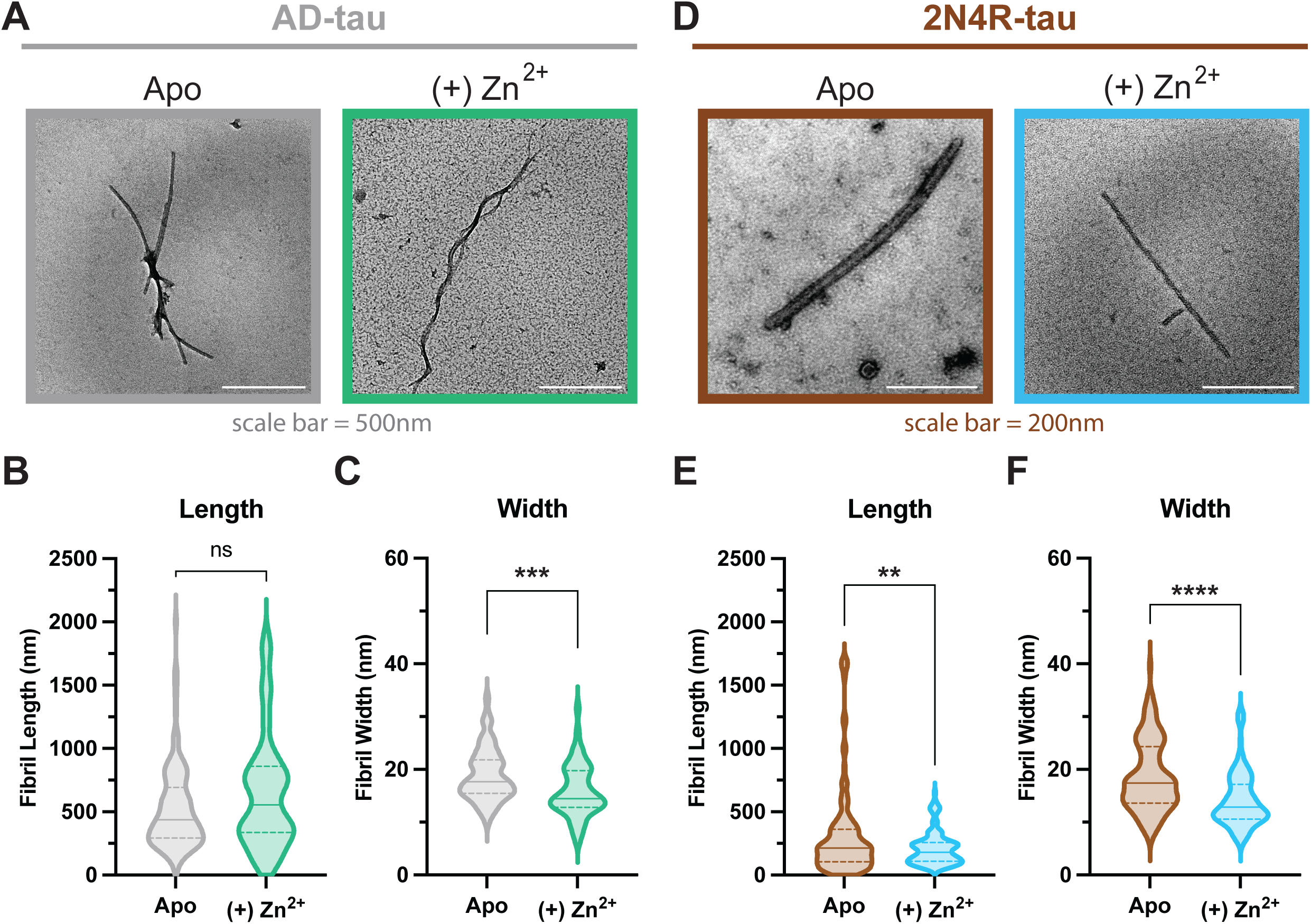
Zinc presence during aggregation impacts fibril morphology for both AD-tau and 2N4R-tau. A) Representative TEM images of AD-tau fibrils aggregated alone ("apo," grey border) or with 1.3 mM ZnSO_4_ ("(+) Zn^2+^," green border). Scale bars = 500 nm. B, C) AD-tau fibril length (B) and width (C). Apo: n = 121 fibrils; (+) Zn^2+^: n = 55 fibrils. Length p = 0.0640 (ns); width ***p = 0.004. D) Representative TEM images of 2N4R-tau fibrils aggregated alone ("Apo," brown border) or with 1.3 mM ZnSO_4_ ("(+) Zn^2+^," blue border). Scale bars = 200 nm. E, F) 2N4R-tau fibril length (E) and width (F). Apo: n = 135 fibrils; (+) Zn^2+^: n = 123 fibrils. Length **p = 0.0020; width ****p < 0.0001. For all figures, violin plots show median (solid line) and upper/lower quartile (dashed lines). Fibrils quantified by a blinded researcher from N = 2 independent aggregation assays; comparisons by Welch’s unpaired t-test.

Knowing that added zinc leads to different tau fibril morphology we also investigated whether or not Zn^2+^ is incorporated within tau fibrils. For this we performed inductively-coupled plasma mass spectrometry (ICP-MS) to determine zinc concentration within the pellet fraction (fibrillar tau) after ultracentrifugation and extensive washing. We found that a significant amount of zinc remained in the pellet fraction of (+) Zn^2+^ conditions for both AD-tau and 2N4R-tau (**Supplemental Figure 3A,B**).

### Fibrils aggregated with Zn^2+^ exhibit altered seeding potential in HEK293T cells

With the knowledge that added zinc impacts aggregation speed and final fibril morphology, we next sought to determine whether fibrils produced in the presence of zinc had altered seeding activity in cells by adapting the previously established Tau-RD FRET Biosensor cells^30^. In this assay, we transfected HEK293T cells with a bi-cistronic plasmid to express two fluorescently tagged constructs of tau that are composed of the microtubule repeat domain of tau (“TauRD”, “K18”) with two point mutations that pre-dispose tau to aggregate (P301L, V337M). These tau fusion proteins (mRuby2-TauRD-LM and mClover2-TauRD-LM) do not aggregate spontaneously in cells, but upon transfection with tau aggregates can be seeded to produce puncta aggregates which can be qualitatively monitored via microscopy and quantitatively measured via FRET flow cytometry (**Supplemental Figure 4A**).

To test the potential for our tau fibrils to seed aggregation in this platform, we pulse-sonicated fibrils to generate seed-competent fragments and transfected equivalent amounts of tau fibril produced with or without Zn^2+^ into our biosensor cells. Both apo and (+) Zn^2+^ AD-tau fibrils were capable of inducing robust puncta in cells as compared to a no seed control (**Figure 4A**). Using flow cytometry to perform quantitative analysis we found the (+) Zn^2+^ seeded condition (green) resulted in a significant increase of cellular aggregation, as measured by FRET-positive events, compared to the apo seed (light grey; **Figure 4B**). As controls we also included conditions where no seed was added (black), where we added ZnSO_4_ alone to cells (purple) or apo seeds with ZnSO_4_ added after fibrillization at the time of seeding (“Apo w/ Zn^2+^”; dark grey).

**Figure 4.**
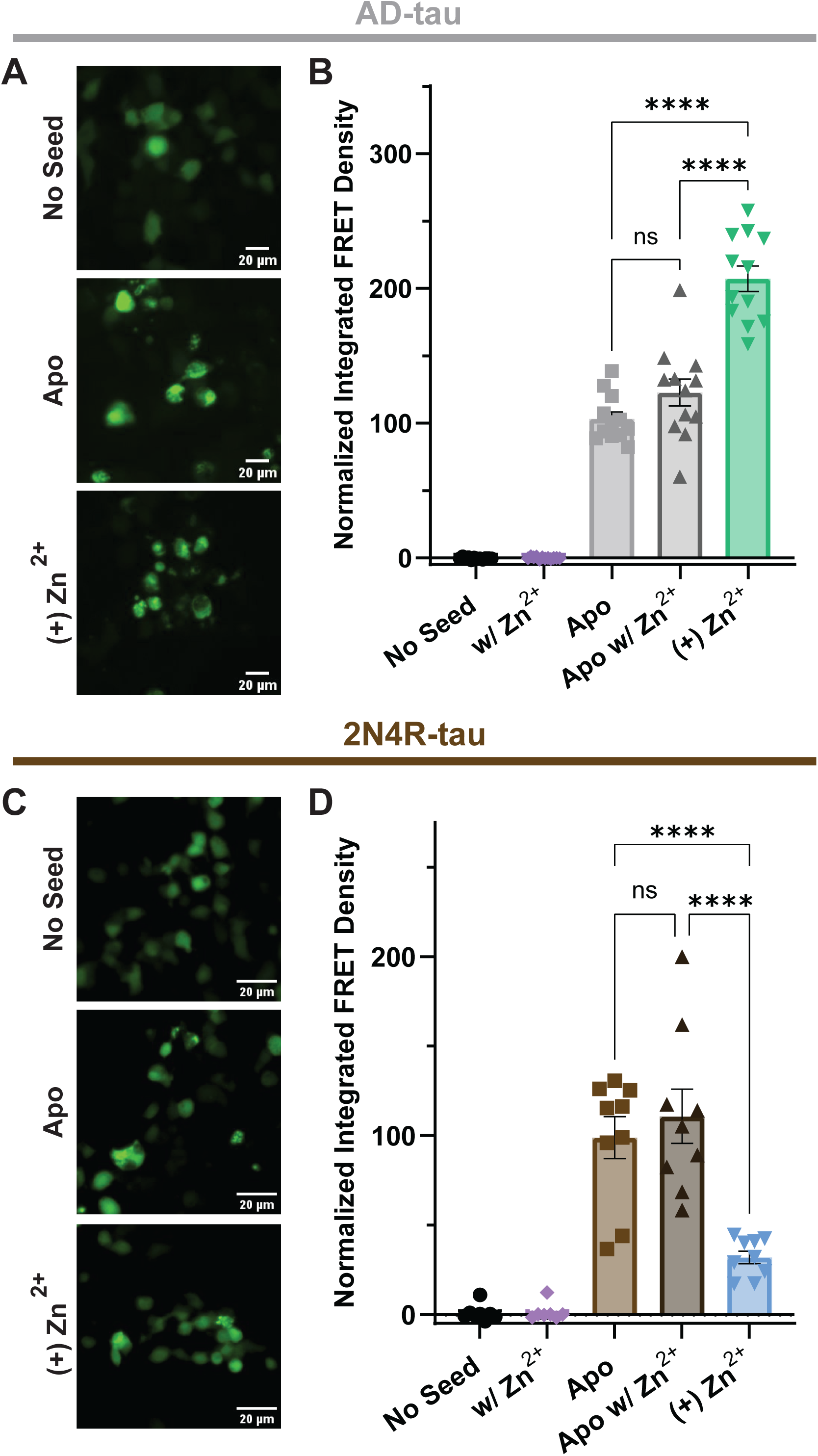
Zinc-made fibrils exhibit altered seeding potential. A) Representative fluorescence images (green channel, 20×) of HEK293T cells expressing a tauRD-FRET biosensor, seeded with apo or (+) Zn^2+^ AD-tau fibrils. Scale bars = 20 µm. B) Normalized integrated FRET density for AD-tau seeds. Conditions: no seed (black), Zn^2+^ alone (purple), apo (light grey), apo + Zn^2+^ post-aggregation (dark grey), (+) Zn^2+^ (green). N = 12 biological replicates; mean ± SEM. Apo vs (+) Zn^2+^, ****p < 0.0001 (one-way ANOVA with Tukey’s post hoc test). C) Representative fluorescence images (green channel, 20×) of HEK293T cells expressing a tauRD-FRET biosensor, seeded with apo or (+) Zn^2+^ 2N4R-tau fibrils. Scale bars = 20 µm. D) Normalized integrated FRET density for 2N4R-tau seeds. Conditions: no seed (black), Zn^2+^ alone (purple), apo (light brown), apo + Zn^2+^ post-aggregation (dark brown), (+) Zn^2+^ (blue). N = 9 biological replicates; mean ± SEM. Apo vs (+) Zn^2+^, ****p < 0.0001 (one-way ANOVA with Tukey’s post hoc test).

For 2N4R-tau fibrils, pulse-sonicated fibrils from both apo and (+) Zn^2+^ fibrils produced visible puncta by microscopy (**Figure 4C**). In contrast to our AD-tau results, we found that (+) Zn^2+^ seeded condition (blue) resulted in decreased aggregation as measured by FRET compared to the apo seed (light brown) and the apo seed with ZnSO_4_ added post-aggregation (“Apo w/ Zn^2+^”, dark brown; **Figure 4D**). Again, we did not detect any aggregation when no seed was added (black) or when Zn^2+^ was added alone to cells (purple). Together, these results demonstrate that the cellular seeding potential is consistent with the phenotypes observed in our *in vitro* aggregation assays and suggest that there may be underlying structural features that contribute to this activity.

## DISCUSSION

Zinc is dysregulated in the brains of Alzheimer’s disease patients, where it is elevated in concentration^11–14^. Yet how zinc shapes tau aggregation, fibril structure, and seeding behavior has remained unclear, in part because prior studies have used varied and often non-comparable experimental conditions. Here, we systematically investigated the impact of Zn^2+^ on tau aggregation using two constructs, the full-length 2N4R-tau isoform and a short AD-tau fragment comprising the ordered core of the AD fibril fold, under identical, heparin-free conditions. Our central finding is that zinc has opposing effects depending on the tau construct used: with AD-tau, zinc accelerates aggregation, produces slightly longer and narrower fibrils, and increases seeding potential, whereas with 2N4R-tau, zinc slows aggregation, produces shorter and narrower fibrils, and decreases seeding potential. These results demonstrate that aggregation kinetics, fibril morphology, and seeding potential are interdependent properties rather than independent outcomes, and they highlight that the choice of tau construct can fundamentally alter the biological conclusions drawn from *in vitro* studies.

The acceleration of AD-tau aggregation by zinc is consistent with prior reports that zinc promotes tau self-assembly. Both tau repeat-domain fragments and full-length tau have been shown to undergo zinc-accelerated aggregation in the presence of heparin^18–20^, and our AD-tau construct results are consistent with these reports. In contrast, the slowing of 2N4R-tau aggregation by zinc conflicts with the acceleration observed for AD-tau and with most prior reports, though there have been bimodal concentration dependence documented in the literature, where higher zinc concentrations shift tau aggregation toward non-fibrillar or inhibited states^19,31^. This construct-dependent divergence likely stems from two factors. First, nearly all prior studies of zinc-tau interactions have been conducted in the presence of heparin, a potent anionic inducer that both accelerates aggregation and produces fibrils with non-disease-relevant structures. We found that heparin inclusion masked the effect of added zinc on both constructs under our conditions (**Supplementary Figure 2**), suggesting that heparin-driven aggregation may override or obscure zinc-dependent modulation. This is consistent with reports that heparin binds divalent cations^32,33^, potentially sequestering zinc and reducing its effective concentration. Second, the N- and C-terminal flanking regions present in 2N4R-tau, but absent in AD-tau, contain additional zinc-binding sites^16,17^, and binding of zinc to these sites may engage intra- or intermolecular interactions that are inhibitory to aggregation rather than promotive. For example, zinc binding to N-terminal histidines could stabilize conformations that disfavor the intermolecular contacts needed for nucleation, or could promote off-pathway species that do not mature into ThT-positive fibrils.

It is also important to acknowledge the broader context of *in vitro* tau aggregation. Full-length tau is notoriously difficult to aggregate without inducers, and several studies have reported that they cannot achieve aggregation of full-length tau in the absence of heparin or other cofactors^34,35^. Our observation that 2N4R-tau aggregates without heparin, albeit slowly and with high day-to-day variability, is itself notable, and the variability we observed (**Supplementary Figure 1**) underscores the challenge of reproducing tau aggregation kinetics across experiments. This variability is a recognized feature of *in vitro* tau aggregation assays and may reflect sensitivity to subtle differences in shaking conditions, temperature, buffer composition and/or protein preparation purity. Additionally, our *in vitro* system lacks the numerous cofactors, lipids, RNA, and post-translational modifications present in the human brain, all of which could modulate zinc’s effects in vivo. The concentration of zinc we used is at the upper end of the physiological range where synaptic zinc can reach ∼1 mM during neurotransmission^24^, but exceeds the average brain zinc concentration of ∼150μM. Future studies should examine dose-dependence, particularly given reports that zinc’s effects on tau can be concentration-dependent and even biphasic^18,19^.

The morphological changes induced by zinc further distinguish the two constructs. AD-tau fibrils formed with zinc were slightly longer and narrower than apo fibrils (**Figure 3A-C**). In contrast, 2N4R-tau fibrils formed with zinc were both shorter and narrower (**Figure 3D-F**) than their apo counterparts. The opposing effect of zinc on fibril length for each construct may contribute to their divergent seeding properties, as longer fibrils could be more fragile and fragment more readily, generating additional seed-competent ends^36^. Both fibril constructs formed with zinc had a reduced width, however all width values fell within the reported range for recombinant tau fibrils by TEM (∼13–20 nm)^37,38^. Direct comparison of our fibril dimensions with published studies is complicated by the near-universal use of heparin as an inducer, which produces fibrils with distinct morphology^29^. Our TEM images do not provide the atomic-resolution structural information that cryo-EM would, and we cannot determine whether the width difference we observe reflects a different protofilament number, a different protofilament fold, or a change in the disordered "fuzzy coat" that surrounds the ordered core. Nevertheless, the reproducible morphological differences between zinc-made and apo fibrils demonstrates that a cofactor can template specific fibril structures under identical protein and buffer conditions. This is significant because cryo-EM structures of patient-derived tau filaments have revealed non-proteinaceous densities enclosed within the filament core^39,40^ as well as additional densities in contact with lysine residues in AD filaments^41^. These densities have been proposed to represent cofactors such as ions, lipids, or hydrophobic moieties, but whether they are passive occupants or active drivers of the resulting fibril fold has remained unclear. Our finding that zinc alone can alter fibril morphology under otherwise identical conditions supports the latter possibility, suggesting that such cofactors may not merely occupy pre-existing cavities but may actively template the fold that encloses them.

The seeding results reveal a striking agreement with the aggregation data: AD-tau fibrils made with zinc seed more efficiently than apo AD-tau fibrils, while 2N4R-tau fibrils made with zinc seed less efficiently than apo 2N4R-tau fibrils. Importantly, the seeding differences are not explained by differences in total fibril mass, as sedimentation assays showed no significant difference in the amount of pelleted tau between zinc and apo conditions for either construct, and equivalent total amounts of seed were transfected. Nor are they explained by zinc itself acting as a seed or directly promoting aggregation in cells, as control conditions with zinc alone or with zinc added post-aggregation did not produce FRET-positive events. Instead, our interpretation is that the seeding differences must arise from structural properties of the fibrils themselves. This finding connects fibril structure to seeding potential, a relationship that has been proposed based on the disease-specificity of tau fibril folds, but has been difficult to demonstrate causally.

Several limitations should be noted. First, our TEM-based morphological analysis do not provide atomic-resolution structural information; cryo-EM will be needed to determine whether zinc alters the protofilament fold, the number of protofilaments, or the inter-protofilament interface. Second, our aggregation assays were performed without post-translational modifications, which are abundant on tau *in vivo* and could modulate zinc binding and aggregation behavior. Third, the zinc concentration used (1 mM) is at the high end of the physiological range, and dose-response studies will be important to determine whether the effects we observe are graded or threshold-dependent. Fourth, the biosensor seeding assay uses a truncated tau construct (TauRD with P301L/V337M mutations) that may not fully recapitulate the seeding behavior of full-length wild-type tau in neurons. Finally, the high day-to-day variability in aggregation kinetics, while consistent with the literature, limits the precision of our kinetic comparisons and underscores the need for standardized aggregation protocols.

This study demonstrates that the trace metal zinc can fundamentally alter tau aggregation kinetics, fibril morphology, and seeding potential, and that these effects are construct-dependent: zinc promotes aggregation and seeding of the AD-tau core fragment but inhibits aggregation and seeding of full-length 2N4R-tau. These findings establish aggregation kinetics, fibril structure, and seeding potential as interdependent properties shaped by cofactor identity, and they argue that the choice of tau construct is not merely a technical detail but a determinant of biological outcome. More broadly, our results support the emerging view that cofactors may be active architects of fibril structure rather than passive occupants, and that understanding cofactor-protein interactions may be key to generating disease-relevant tau fibrils *in vitro*.

## EXPERIMENTAL PROCEDURES

### Molecular Cloning

The AD-tau construct (residues G304-E380 of 2N4R) was cloned into the pRK172 plasmid backbone using Gibson assembly. The correct DNA sequence was confirmed through full plasmid sequencing (Plasmidsaurus).

### Recombinant Tau Purification

AD-tau or 2N4R-tau were purified as previously reported^42,43^. Briefly, the tau construct plasmid was transformed into *E. coli* BL21 cells (DE3) and grown overnight in LB with ampicillin, shaking at 200rpm at 37°C. 25mL of these pre-cultures were then used to inoculate 1L of TB (with ampicillin) and left to shake at 140rpm at 37°C until OD_600_ = 0.8 is reached. 100mL of NaCl/betaine solution (5M NaCl:5M at 50:1 ratio) is added, left to continue shaking for 30 minutes, now at 30°C. Cells are induced with 500µM IPTG and left to shake overnight at 18°C. Cells were centrifuged to obtain a cell pellet and the pellet was resuspended in lysis buffer (50mM MES pH 6.5, 5mM DTT, 1mM PMSF, 1mM EGTA + cOmplete protease inhibitor tablets (Roche)), dounce homogenized, and sonicated on ice for a total of 10 minutes, pulsing on for 30 seconds, off for 30 seconds at 30% amplitude. NaCl was added to lysate to obtain a final concentration of 500mM then boiled for 15 minutes in a water bath followed by centrifugation at 20,000xg for 30 minutes at 4°C. Supernatant was precipitated with ammonium sulfate (30% m/v) on ice for 15 minutes and centrifuged at 20,000xg for 30 minutes at 4°C. Pellet was collected, resuspended in Buffer A (20mM MES pH 6.0, 50mM NaCl, 1mM EGTA, 0.2mM MgCl2, and 2mM DTT) and dialyzed overnight in Buffer A at 4°C.

Dialyzed solution was filtered through a 0.45µm filter and subjected to cation exchange chromatography with a HiPrep SP HP 16/10 column equilibrated in Buffer A. Protein was eluted from the column with a linear gradient of Buffer B (20mM MES pH 6.0, 1M NaCl, 1mM EGTA, 0.2mM MgCl2, and 2mM DTT) and purified protein was visualized on an SDS-PAGE gel. Fractions were combined, concentrated, and buffer exchanged into 1x Phosphate Buffered Saline, pH 7.4 (Gibco). The final concentration of the purified protein was determined using a Pierce BCA Protein Assay Kit (ThermoFisher). Purified protein was divided into 100µL aliquots, flash frozen in LN_2_, and stored at −80°C until needed.

### In vitro Thioflavin T Tau Aggregation Assay

Reactions were prepared for each condition such that final concentrations were 50µM tau, 2mM TCEP, 5mM MgCl_2_, and 10mM KCl, unless stated otherwise in the text. Some reactions included 20µM Thioflavin T (ThT), and/or 12.5µM heparin and/or 1.3mM ZnSO_4_, as indicated in the text. Heparin (Sigma; 15-19kDa), Tris(2- carboxyethyl)phosphine (TCEP) (Sigma), and ZnSO_4_ heptahydrate (Sigma) were made fresh immediately prior to use. “Blank” wells, with all components excluding tau protein, were run in triplicate as a background fluorescence control. Reactions were loaded into a 96-well half-area black plate (Greiner Bio) containing two polytetrafluoroethylene (PTFE) beads, and the plate was sealed with an opti-clear film to prevent evaporation. The plate was placed into an Agilent BioTek Synergy H1 Microplate Reader programmed with continuous orbital shaking at 425cpm. A 444nm excitation wavelength and 485nm emission wavelength were used to record ThT fluorescence over time. Wells without ThT were used for FRET cellular seeding assays. All samples were collected, ultracentrifuged (see “Isolation of Tau Fibrils”) and flash frozen with LN_2_ and stored at −80°C for future use.

ThT fluorescence data was collected from the Agilent Plate reader software and analyzed using GraphPad PRISM Version 11. “Blank” wells were subtracted from the raw values of all experimental condition wells. Readings from time points after ThT fluorescence plateaued were omitted, and each technical replicate was normalized using the average of the first 10 readings for all wells of that condition as 0% and the average of the last 10 time points for all wells of that condition at plateau as 100%. A nonlinear regression was performed on the normalized data using the Goempertz’s growth function: 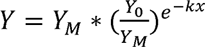

Where *Y_O_* is the minimum normalized relative fluorescence, *Y_M_* is the maximum normalized relative fluorescence, *x* is time, and *t* is the constant determined during fitting which is in hours^-1^. *Y_O_,Y_M_,* and *k*, are reported as well as 50, the time, in hours, it took for ThT fluorescence to reach 50% is calculated.

### Isolation of Tau Fibrils

Following aggregation, replicate wells were combined and ultracentrifuged at 117,000xg for 80 minutes at 4°C. The supernatant fraction, containing monomeric tau, was removed and pellet was washed with 100uL PBS pH 7.4. The pellet fraction, containing fibrillar tau, was resuspended in PBS pH 7.4. The tau concentration was determined by BCA and samples are flash frozen in LN_2_ and stored at −80°C for further use

### Determining Ratio of Supernatant:Pellet Fractions by SDS-PAGE

Supernatant and pellet fractions were heated at 95°C for 10 minutes in 1xSDS Loading Dye (62.5 mM Tris-HCl (pH 6.8), 2% (w/v) Sodium dodecyl sulfate (SDS), 10% glycerol, 0.01% (w/v) bromophenol blue, 50mM Dithiothreitol (DTT)). Samples were run on 10-20% SDS-PAGE gels, stained with Imperial Protein Stain (ThermoFisher) and imaged using the BioRad ChemiDoc Go Imaging System. Analysis of band intensity was performed with Fiji ImageJ software.

### HEK293T Tau-FRET Seeding Assay

HEK293T were plated at 70,000 cells/well in a 24-well plate and incubated overnight at 37°C, 5% CO_2_. Cells were transfected with Lipofectamine 2000 Transfection reagent to introduce the Tau-RD FRET construct (mRuby-TauRDP301L,V337M-P2A-mClover-TauRDP301L,V337M). Plates were incubated overnight at 37°C, 5% CO2 and inspected for green fluorescence to ensure proper transfection. Cells were seeded with tau fibrils (500nM, unless otherwise stated) that were briefly pulse sonicated prior to use (pulse for 20 seconds on, 20 seconds off, at 30% amplitude for a total sonication time of 5 minutes). Fibrils were transfected into cells using Lipofectamine 2000 according to the standard protocol for DNA transfection. Control wells (“no seed”) were seeded with Lipofectamine 2000 alone. Plates were incubated for 48 hours at 37°C, 5% CO_2_. Media in each well was aspirated, washed with PBS pH 7.4 and lifted with trypsin dilute 1 to 4 in PBS. Trypsin was inactivated with FBS (10%). Cells were gently resuspended in PBS, pH 7.4, transferred to a 96-well plate and run on an Attune NxT Flow Cytometer (ThermoFisher Scientific). Each sample was analyzed for 10,000 single cell events. Gating was conducted as follows: all events were gated to identify HEK293T cells, then those cells were gated to identified single cells. Single cells were gated by the detection of mClover2 emission to identify transfected cells, and transfected cells were gated by detection of mRuby2 emission to identify FRET positive cells (**Supplemental Figure 4**). The Integrated FRET Density for each sample was calculated by multiplying the %FRET positive cells in each well by the median mRuby2 fluorescence in that well. Within each independent trial, integrated FRET density values were normalized by setting the mean of the no-seed condition to 0% and the mean of the apo seeded condition to 100%, using the same scaling factor for all conditions within that trial, to control for day-to-day variation in transfection efficiency.

### Zinc detection with Inductively Coupled Plasma Mass Spectrometry (ICP-MS)

Tau fibrils (pellet fractions) are diluted 1:9 (for a total volume of 1mL) and given to the UMass Mass Spectrometry Core to conduct ICP-MS and determine the level of zinc present in each sample.

### Transmission Electron Microscopy

300-mesh carbon-coated copper grids (Electron Microscopy Sciences) were glow-discharged using Pelco EasiGlow machine for 30 seconds at 15mAmps. 5µL of aggregated tau samples were added to the grid, incubated for 90 seconds, blotted and incubated for an additional minute. Samples were stained with 1% uranyl acetate and stored until imaging.

Grids were imaged on the FEI Tecnai T12 microscope at 120kV high tension, with at least 10 squares of each grid imaged. The images were analyzed by a researcher blinded to the identity/condition using Fiji ImageJ software measurement tool to quantify fibril length and width.

### Statistical Analysis

Data analysis was performed using GraphPad Prism Software (version 11.0.2). Unless otherwise noted, graphs show mean values, with error bars representing SEM. Statistical significance of differences between conditions was determined in a few different ways, depending on the data.

For data comparing only 2 conditions, t-tests were conducted. Paired t-tests were performed most often, as most experiments (ThT aggregation, ratio of supernatant to pellet, ICP-MS) were conducted with the apo and (+) Zn^2+^ treated as a pair. Welch’s unpaired t-tests were performed when comparing fibril lengths and widths (TEM) as numerous individual fibrils were quantified.

For data comparing more than 2 conditions (cellular FRET seeding assays), one-way ANOVAs were performed with multiple comparisons using Tukey’s post-hoc test.

## ACKNOWLEDGEMENTS

This work was supported by grants from the National Institutes of Health (R00 AG064116 and R01 AG077672) to J.N.R. This work was partially supported by a National Research Service Award T32 GM139789 from the National Institutes of Health award to E.L.P. We thank Dr. Steve Eyles (UMass Amherst) for his assistance with the ICP-MS. We are thankful to the University of Massachusetts Core Facilities for use of the Mass Spectrometry Core and Electron Microscopy Core. Some figure images were created using Biorender.

## SUPPLEMENTAL FIGURE LEGENDS

**Supplemental Figure 1.** Biological replicates of AD-tau and 2N4R-tau ThT aggregation and comparison of shaking conditions

A) ThT aggregation curves for N = 7 AD-tau biological replicates (single orbital shaking), each with n = 3–6 technical replicates. 50 µM tau alone ("Apo," grey) or with 1.3 mM ZnSO_4_ ("(+) Zn^2+^," green). Each technical replicate normalized individually; mean ± SEM plotted per biological replicate.

B) ThT aggregation curves for N = 6 2N4R-tau biological replicates ("Apo," brown; "(+) Zn^2+^," blue). Three assays with single orbital shaking, three with double orbital.

C,D) Y_50_ (hours) compared between single and double orbital shaking for apo (C) and (+) Zn^2+^

(D). Each shaking type: N = 3 biological replicates with n = 3–6 technical replicates each. Each point represents the mean of one biological replicate. No significant difference between shaking types for either condition (paired t-tests).

**Supplemental Figure 2.** H**e**parin **inclusion masks the effect of Zn^2+^ on aggregation kinetics and seeding potential**

A) Representative AD-tau ThT aggregation curve with added heparin, single orbital shaking. 50µM tau + 12.5µM heparin ("Apo +hep," grey) or 50µM tau + 12.5µM heparin + 1.3 mM ZnSO_4_ ("(+) Zn^2+^ +hep," green). n = 3 technical replicates, each normalized individually; mean ± SEM.

B) Representative 2N4R-tau ThT aggregation curve with added heparin, single orbital shaking. 50µM tau + 12.5µM heparin ("Apo +hep," brown) or 50µM tau + 12.5µM heparin + 1.3 mM ZnSO_4_ ("(+) Zn^2+^ +hep," blue). n = 3 technical replicates, each normalized individually; mean ± SEM.

C) Y_50_ (hours) for AD-tau + heparin. N = 5 biological replicates, each with n ≥ 3 technical replicates. Each point represents the mean of one biological replicate. No significant difference (paired t-test).

D) Y_50_ (hours) for 2N4R-tau + heparin. N = 3 biological replicates, each with n ≥ 3 technical replicates. Each point represents the mean of one biological replicate. No significant difference (paired t-test).

E) AD-tau pellet fraction as a percentage of total tau (Sup + Pel), quantified by densitometry, from 3 biological replicates. Mean ± SEM plotted. No significant difference using a paired t-test.

F) 2N4R-tau pellet fraction as a percentage of total tau (Sup + Pel), quantified by densitometry, from 2 biological replicates. Mean ± SEM plotted. No significant difference using a paired t-test.

G) Representative SDS-PAGE of supernatant (Sup) and pellet (Pel) fractions after ultracentrifugation of AD-tau aggregation.

H) Representative SDS-PAGE of supernatant (Sup) and pellet (Pel) fractions after ultracentrifugation of 2N4R-tau aggregation.

**Supplemental Figure 3: Zinc is incorporated into both AD-tau and 2N4R-tau fibrils**

A) Zinc (ppb) per µmol AD-tau fibrils, measured by ICP-MS. N = 3 biological replicates; mean ± SEM.

B) Zinc (ppb) per µmol 2N4R-tau fibrils, measured by ICP-MS. N = 2 biological replicates; mean ± SEM.

**Supplemental Figure 4. Flow cytometry gating scheme and independent trials of the cellular seeding assay**

A) Gating scheme for flow cytometry analysis of the FRET seeding assay. No-seed and seeded "FRET positive" gates are shown to illustrate the difference between conditions.

B) Integrated FRET density for 4 independent trials of AD-tau seeding. Conditions: no seed (black), Zn^2+^ alone (purple), apo (grey), apo + Zn^2+^ post-aggregation (dark grey), (+) Zn^2+^ (green). Each trial: n = 3 biological replicates (total N = 12); mean ± SEM. *p < 0.0332, **p < 0.0021, ***p < 0.0002, ****p < 0.0001 (one-way ANOVA with Tukey’s post hoc test).

C) Integrated FRET density for 3 independent trials of 2N4R-tau seeding. Conditions: no seed (black), Zn^2+^ alone (purple), apo (brown), apo + Zn^2+^ post-aggregation (dark brown), (+) Zn^2+^ (blue). Each trial: n = 3 biological replicates (total N = 9); mean ± SEM. *p < 0.0332, **p < 0.0021, ***p < 0.0002, ****p < 0.0001 (one-way ANOVA with Tukey’s post hoc test).

