## Supplementary figures and images for "Zinc Differentially Modulates Tau Aggregation, Fibril Morphology, and Prion-like Seeding in a Construct-Dependent Manner"

### Supplemental Figure 1

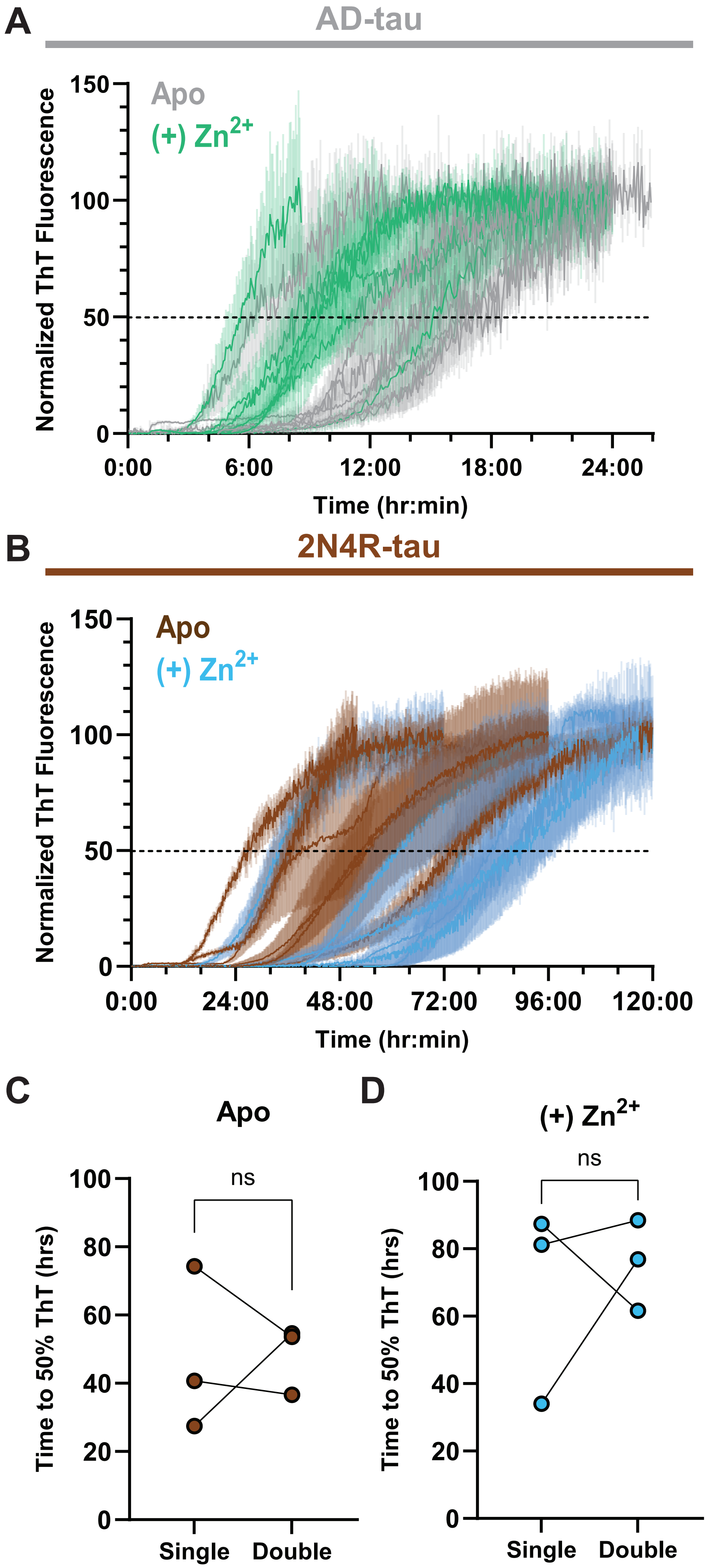

### Supplemental Figure 2

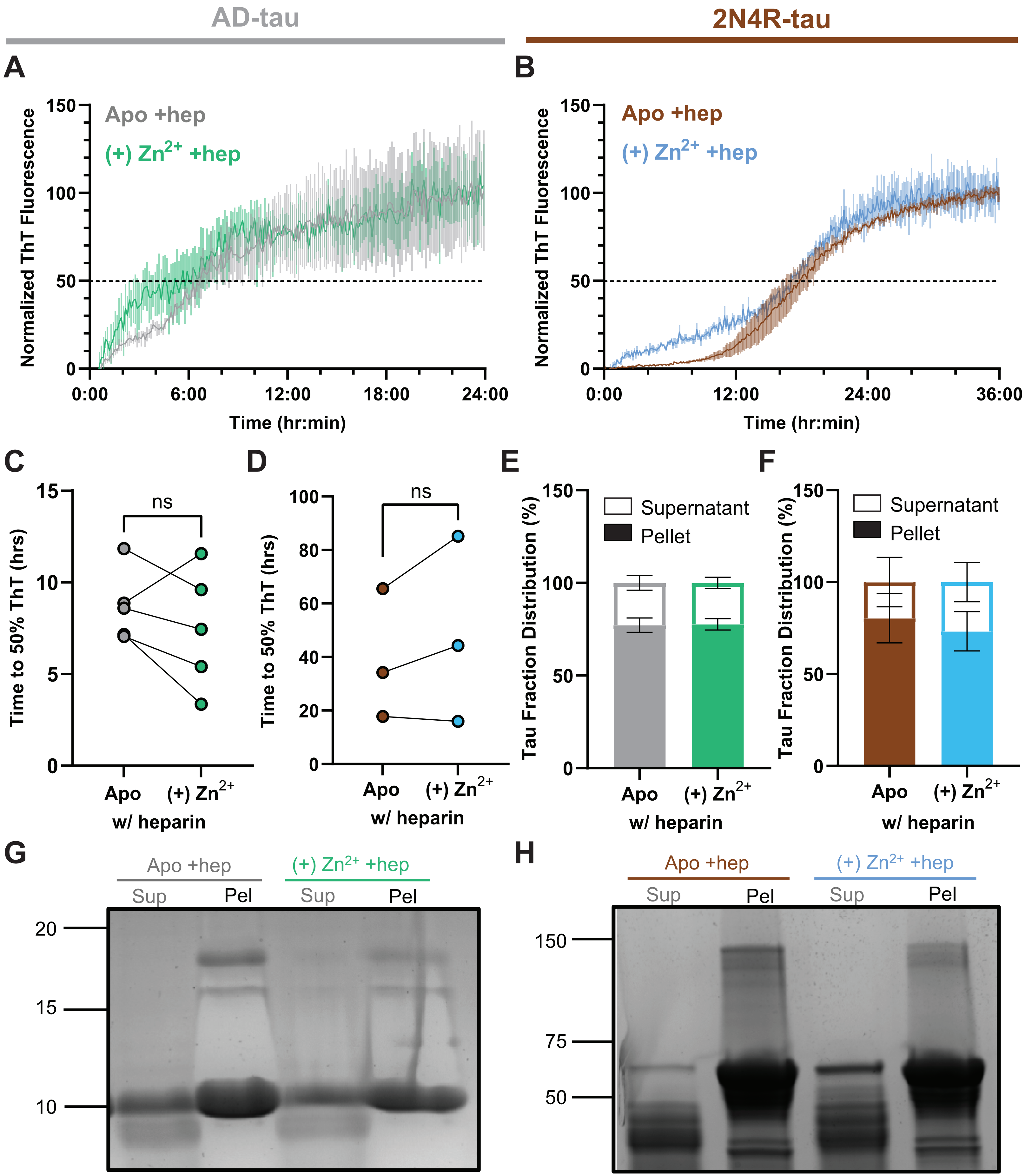

### Supplemental Figure 3

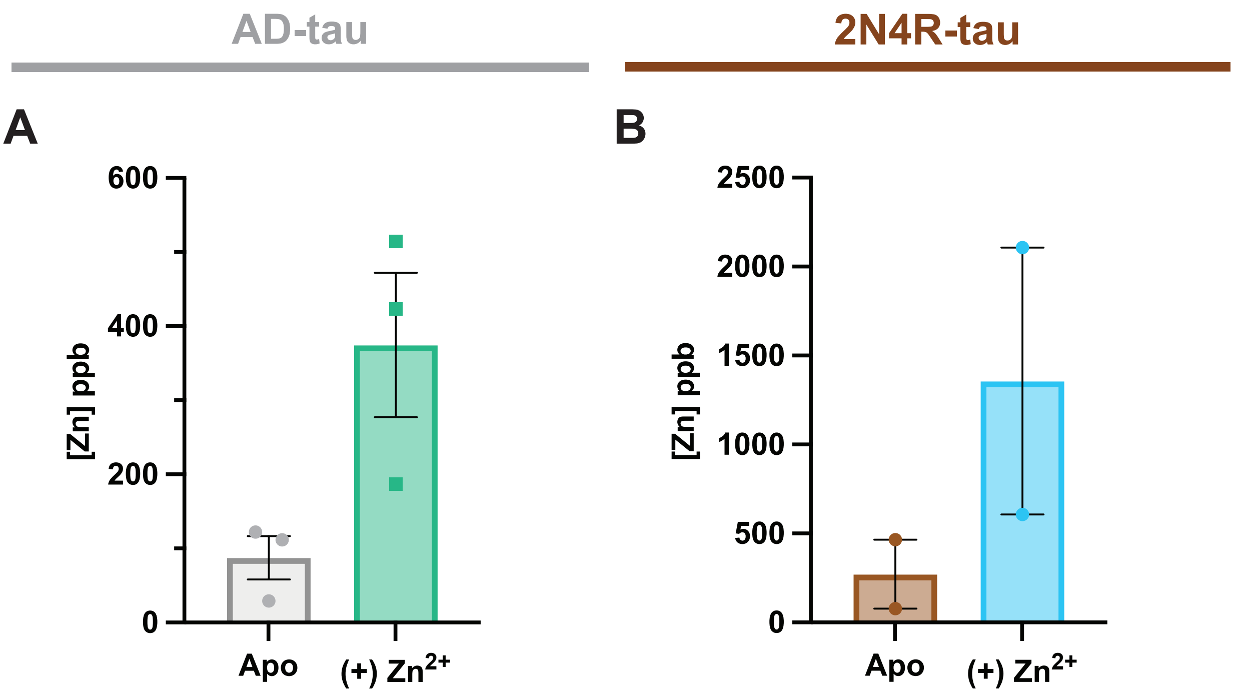

### Supplemental Figure 4

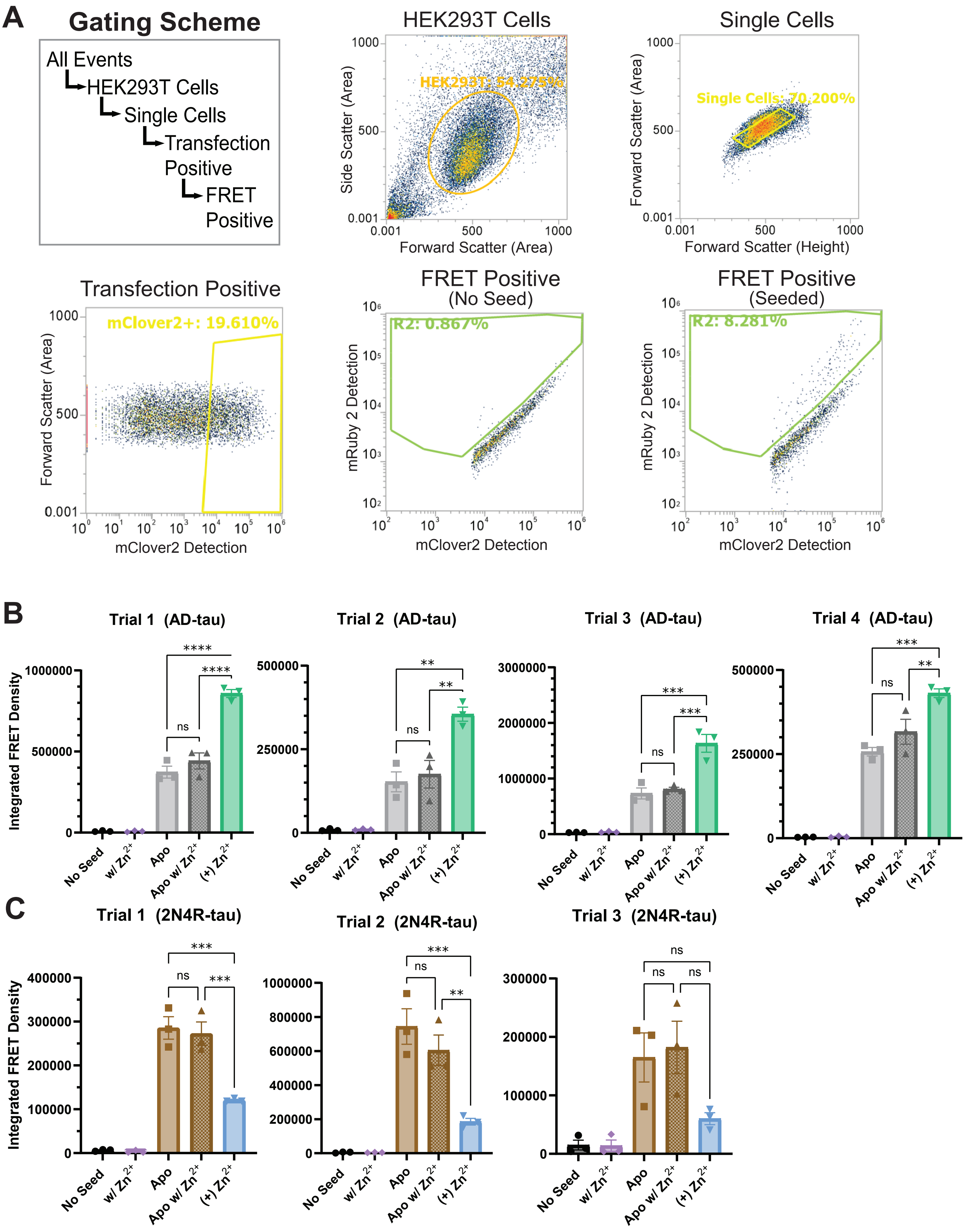
